# Grazing pressure-induced shift in planktonic bacterial communities with the dominance of acIII-A1 actinobacterial lineage in soda pans

**DOI:** 10.1101/2020.06.04.133827

**Authors:** Attila Szabó, Kristóf Korponai, Boglárka Somogyi, Balázs Vajna, Lajos Vörös, Zsófia Horváth, Emil Boros, Nóra Szabó-Tugyi, Károly Márialigeti, Tamás Felföldi

## Abstract

Astatic soda pans of the Pannonian Steppe are unique environments with respect to their multiple extreme physical and chemical characteristics (high daily water temperature fluctuation, high turbidity, alkaline pH, salinity, polyhumic organic carbon concentration, hypertrophic state and special ionic composition). However, little is known about the seasonal dynamics of the bacterial communities inhabiting these lakes and the role of environmental factors that have the main impact on their structure. Therefore, two soda pans were sampled monthly between April 2013 and July 2014 to reveal changes in the planktonic community. By late spring in both years, a sudden shift in the community structure was observed, the previous algae-associated bacterial communities had collapsed, resulting the highest ratio of actinobacteria within the bacterioplankton (89%, with the dominance of acIII-A1 lineage) ever reported in the literature. Before these peaks, an extremely high abundance (>10,000 individuum l^−1^) of microcrustaceans (*Moina* and *Arctodiaptomus*) was observed. OTU-based statistical approaches showed that in addition to algal blooms and water-level fluctuations, zooplankton densities had the strongest effect on the composition of bacterial communities. In these extreme environments, this implies a surprisingly strong, community-shaping top-down role of microcrustacean grazers.

## Introduction

The Pannonian Steppe is a part of the Eurasian steppe; it is situated within the Carpathian Basin, surrounded by the Carpathian Mountains. Soda lakes or smaller pans, as in elsewhere in the steppe, are integral parts of the landscape. They are particularly numerous within the Carpathian Basin, which is their most western occurrence in Eurasia (Boros and Kolpakova, 2018). Soda pans in this region represent multiple extreme environments with respect to their special ion composition, alkaline pH, salinity, high turbidity, humic material and nutrient content, and special underwater light-limited conditions (Boros *et al.*, 2014, 2017, 2020). The lakes can be divided into two ecologically distinct types: some of them are ‘turbid’ because of the immense number of inorganic suspended particles, and others are ‘colored’ due to the high concentration of dissolved humic matter (Boros *et al.*, 2014). Planktonic algal blooms occur frequently in these lakes, since the high nutrient content (inorganic nitrogen and phosphorus) is present throughout the year, mainly due to the guano of migratory birds and decaying plant material originating from the shoreline vegetation (Somogyi *et al.*, 2009; Boros *et al.*, 2016). During summer months, filamentous cyanobacteria bloom in the shallow waters, while at temperatures under 15°C, pico-sized green algae become abundant and may form visible algal blooms (Somogyi *et al.*, 2009; Pálffy *et al.*, 2014). Fish are absent from these pans due to their intermittent character, and partly because of this microcrustaceans, especially the natronophilic *Arctodiaptomus spinosus* (Copepoda: Calanoida) and *Moina brachiata* (Cladocera) can be abundant by late spring (Horváth *et al.*, 2014; Tóth *et al.*, 2014). Due to the complexity of aquatic food webs and bottom-up effects, only a few reports were published on how zooplankton regulate natural aquatic microbial communities (e.g., Eckert *et al.*, 2013). Since the microbial communities of these pans are not limited by nutrients, these environments pose as natural field models to study the grazing effect of zooplankton (rotifers and microcrustaceans) on bacterioplankton composition.

Prokaryotic communities of many alkaline lakes were investigated in detail (e.g., Antony *et al.*, 2013; Lanzén *et al.*, 2013; Vavourakis *et al.*, 2016; Andreote *et al.*, 2018; Edwardson and Hollibaugh, 2018, Zorz *et al.*, 2019), but no comprehensive study focusing on the seasonal dynamics and biological relationships of soda lake planktonic microbial communities has been performed so far. Nevertheless, soda lakes can be found on almost every continent, and these extreme environments represent a significant fraction of continental saline aquatic habitats and an important source of unexplored microbial biodiversity (Hammer 1986, Grant 2004, Sorokin et al. 2014).

The relatively easily accessible pans of the Pannonian Steppe provide a unique opportunity to reveal temporal patterns in the microbial community structure, as was reported previously in the case of freshwater and marine communities (see, e.g., Eiler *et al.*, 2012; Fuhrman *et al.*, 2015). Therefore, monthly sampling was carried out for more than a year in two *ex lege* protected soda pans representing different types, the colored Sós-ér pan and the turbid Zab-szék pan situated in the Kiskunság region (Hungary). In order to put these findings into a broader ecological context, microscopic and zoological investigations were carried out in parallel.

## Results

### Environmental conditions

The water depth of the pans changed remarkably throughout the study period (in the turbid Zab-szék pan: 2-45 cm, in the colored Sós-ér pan: 0-50 cm). The water level in the turbid pan was under 12 cm from February to July 2014 and under 3 cm from April to June 2014. Salinity values also varied greatly, from subsaline (1.5 - 2.4 g l^−1^, associated with high water level during the spring of 2013) to almost mesosaline (18.9 g l^−1^, close to desiccation). The pH was alkaline throughout the study period (turbid pan: 9.1 - 10.2; colored pan: 8.3 - 10.0) and was the lowest in the colored pan during the spring of 2013. The water temperature remained below 16°C from September to March, with the lowest value measured in December 2013. During the summer months, the water temperature varied between 25°C and 31°C. The total organic carbon content was generally higher in the colored pan [the highest values (855, 1223 mg l^−1^) were observed before desiccation]. In the colored pan, organic carbon was generally present in dissolved form, while in the turbid pan, it was composed of both particulate and dissolved matter. The inorganic suspended solids (ISS) content was much higher on average in the turbid pan than in the colored pan. Prokaryotic cell numbers varied between 1.42 × 10^7^ and 4.74 × 10^8^ cells ml^−1^. The environmental parameters measured are given in Supplementary Table S1.

### Phytoplankton

Based on their chlorophyll *a* content, pans were hypertrophic; a winter/spring algal bloom was observed from November 2013 to April 2014. During the algal bloom, nanoplanktonic and picoeukaryotic algae dominated in the colored pan, while in the turbid pan an exclusive picoeukaryotic dominance was found (Supplementary Table S1 and S3). Moreover, in the turbid pan, the phytoplankton was dominated by picoalgae through the entire study period. In August 2013, picocyanobacteria (genus *Synechococcus*, Supplementary Fig. S2) proliferated massively, then in October the pico-sized green algae (*Chloroparva*, *Choricystis* and other trebouxiophyceaen green algae, Supplementary Fig. S3) became dominant; their abundance peaked in February 2014 (with 3 × 10^5^ μg l^−1^ biomass). Picoalgal cell numbers were higher by one order of magnitude there than in the colored pan. Nanoplanktonic algae were detected only on three sampling dates: *Nitzschia palea* and *Chlamydomonas* sp. in August 2013, *Nitzschia palea* in December 2013 and *Cryptomonas* sp. in March 2014, the latter with a high biomass (3,450 μg l^−1^).

Contrary to in the turbid pan, larger nanoplanktonic algae dominated in the colored pan. During the spring and early summer of 2013, algal cells were scarce and mostly nanoflagellates (*Cryptomonas*, *Rhodomonas*, *Chlamydomonas*) were present (chlorophyll *a* concentration 0.5-3.6 μg l^−1^). In July, only diatoms were detected (*Nitzschia*, *Cocconeis* and *Cymbella* species). Before the desiccation of the pan, cyanobacteria (*Anabaenopsis* sp.), pico-sized green algae, euglenoids and green flagellated algae (*Chlamydomonas* sp.) were abundant in the water (the chlorophyll *a* concentration was 460 μg l^−1^ in August 2013). After refilling in November 2013, the chlorophyll *a* concentration values were high (between 350 and 380 μg l^−1^) until February 2014, and pico-sized green algae, euglenoids (*Euglena* and *Phacus*) and green flagellated algae (*Carteria* and *Chlamydomonas* species) dominated. Before the desiccation in July 2014, *Navicula* sp. was also present. In December 2013, the filamentous cyanobacterium *Oscillatoria* was also detected.

### Zooplankton

In the turbid pan, cladocerans and copepods peaked in May 2014, with a surprisingly high abundance of 14,160 and 17,760 individuum l^−1^ (Supplementary Table S1). These groups were present throughout the year, with an average abundance of 143 and 636 individuum l^−1^, respectively. The average abundance of rotifers in this pan was 637 individuum l^−1^, and they peaked in September 2013, with 5148 individuum l^−1^. In the colored pan, the abundance of small crustaceans was an order of magnitude lower, with maximum values in July 2013 and in March 2014. Rotifer numbers were high in May 2013, between July and August 2013, before the desiccation of the pan.

### Diversity and composition of the prokaryotic community

Based on the qPCR results, Archaea represented only a minor fraction (13% on average) of the prokaryotic community (Supplementary Fig. S4); however, they were detected in most of the samples. Their average proportion of the community was higher in the turbid pan (19%), and their highest relative abundance (47%) was also detected in the turbid pan in the sample taken in June 2014.

The number of bacterial OTUs, the estimated richness and the diversity indices varied across the sampling period (Supplementary Fig. S5). Richness and diversity values were generally higher in the turbid pan. In this pan, the lowest values were detected in July 2013 and June 2014, during the actinobacterial relative abundance peaks. The highest richness values from this pan were obtained from September to October 2013. In the colored pan samples, the lowest richness was detected in the April 2013 sample, while the highest richness was estimated in the March 2014 sample. Diversity estimators in the turbid pan were lowest in June 2014 and highest in May and December 2013, and in March 2014. In the colored pan, the lowest values were recorded in April 2013 and the highest in June 2014. Based on the two-way PERMANOVA test, the bacterial communities of the pans and seasons were significantly different (p < 0.001) in the study period. From the 3127 OTUs obtained, 532 were shared between both pans, 1858 were detected only from the turbid pan and 737 only from the colored pan. However, the vast majority of the sequences (81%) belonged to the shared OTUs, and thus the dominant OTUs were the same in both pans. Based on the Bray-Curtis similarity index, the communities showed similar seasonal dynamics (Supplementary Fig. S6). Changes in the taxonomic compositions of the pans had the same tendency from April to May 2013, and the same pattern was observed between February and May 2014 in the turbid pan and between March and June in the colored pan. This common phenomenon was the result of remarkable changes in the relative abundance of actinobacteria (Fig. 1). Similarities in the bacterial community structures were visualized by NMDS ordination based on a Bray-Curtis distance matrix (Fig. 2). The following environmental variables were fitted significantly (p ≤ 0.05) onto the NMDS ordination: depth, pH, salinity, concentration of chlorophyll *a*, dissolved organic carbon (DOC), particulate organic carbon (POC), ISS, picocyanobacterial and picoeukaryotic algal biomass, copepod and cladoceran abundance. Based on the SIMPER test, the following OTUs were responsible for the at least 40% differences among the samples: OTU1 (acIII-A1), OTU15 (Cyclobacteriaceae), OTU10 (*Flavobacterium*), OTU9 (*Flavobacterium)*, OTU3 (*Nitriliruptor*), OTU2 (*Synechococcus)*, OTU6 (bacII), OTU7 (*Rhodobaca*), OTU8 (*Hydrogenphaga)*, OTU4 (*Ilumatobacter*), OTU5 (acTH1), OTU14 (Actinobacteria), OTU13 (*Luteolibacter*), OTU11 (acSTL), OTU31 (*Algoriphagus*), OTU29 (*Limnohabitans*), OTU16 (alfVI) and OTU27 (alfVI) (OTU serial numbers correspond to their abundance in the whole dataset; lower numbers with a higher constitution). Most of the taxa were related to higher algal densities, while betaproteobacteria were related to the high water level and low salinity period of the pans (during the spring of 2013). These samples (especially from the colored pan) were the most distinct from the others. An inverse relationship was observed between prokaryotic cell numbers and depth. Samples from the low water-level period were fitted with pH, salinity, the concentration of DOC, POC and ISS, because of the evaporation and concentration of lake water. Chlorophyll *a* concentration, nanoplankton biomass and an *Euglena* chloroplast OTU (colored pan), picoeukaryotic algal biomass (turbid pan) fitted with samples from the winter algal bloom period. Picocyanobacterial biomass fitted with the summer samples of the turbid pan. The abundance of microcrustaceans fitted with the samples having high relative abundance of actinobacteria, except the June 2014 sample from the turbid pan. Based on the significantly fitted environmental parameters and the NMDS ordination, water regime (depth and salinity), phytoplankton (algal biomass and composition) and microcrustaceans (their abundance) shaped the planktonic bacterial community structure.

**Fig. 1.**
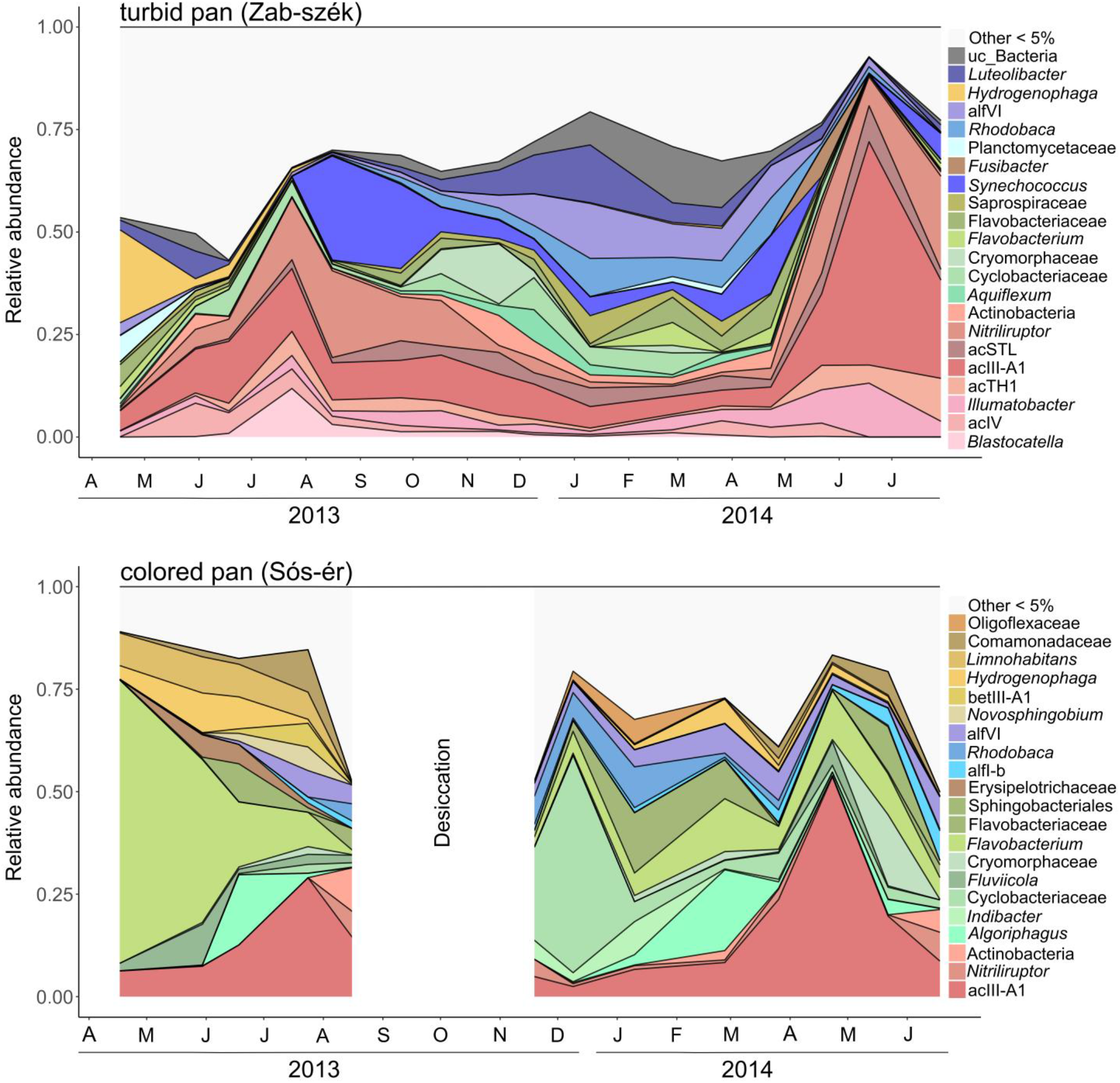
Seasonal changes in the bacterial OTU composition of the turbid and colored pan between April 2013 and July 2014. ‘Other’ represented taxa has less than 5% relative abundance. Letters on the x-axis are the abbreviations for months. Abbreviation: uc_Bacteria (unclassified Bacteria).

**Fig. 2.**
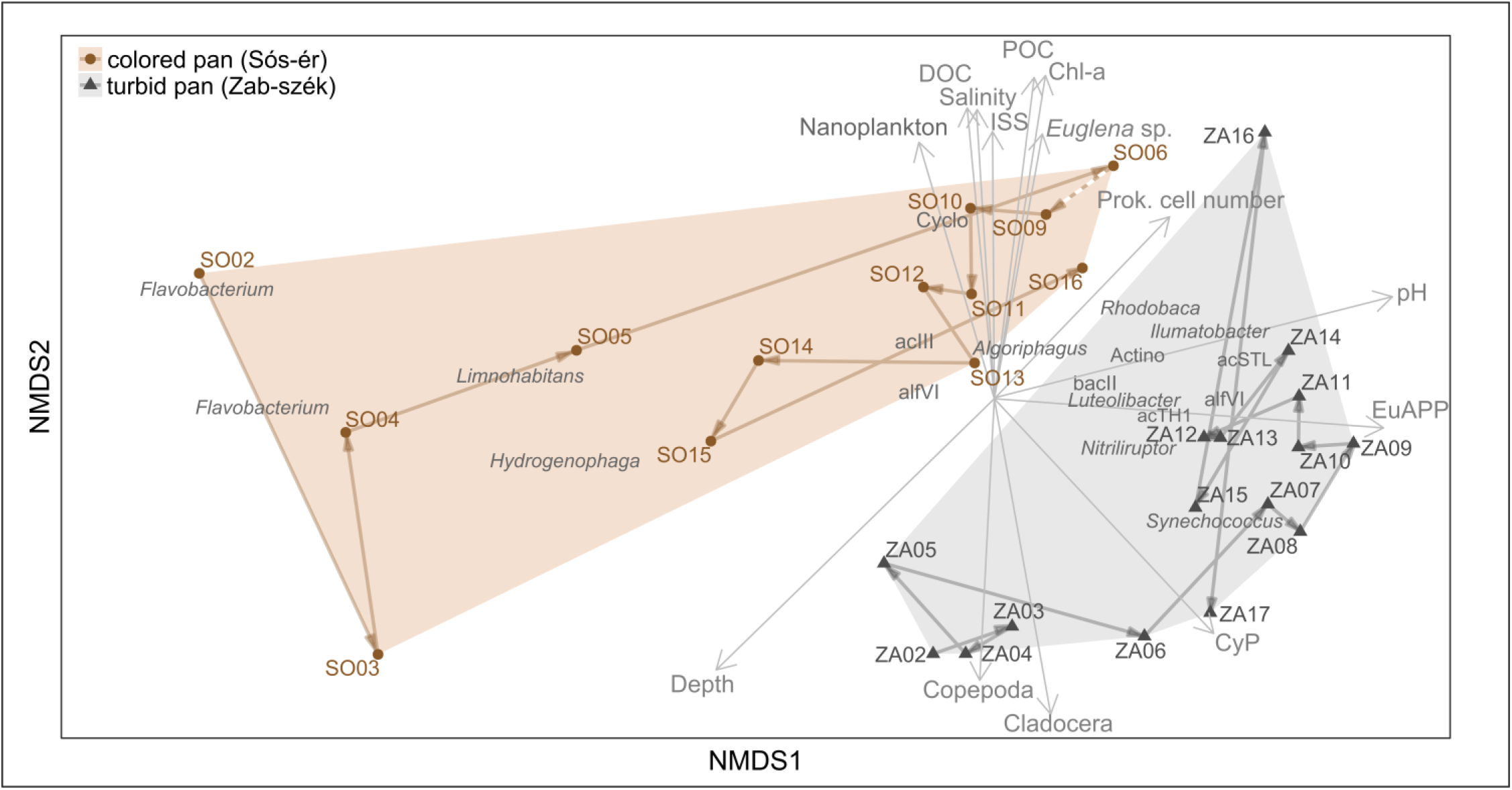
NMDS ordination of soda pan samples collected between April 2013 and July 2014 based on the Bray-Curtis distance of the bacterial OTUs (stress: 0.15). Based on SIMPER analysis, OTUs responsible for 40% dissimilarity among samples are shown in gray. Significantly fitted (p < 0.05) environmental variables are shown with gray vectors. Abbreviations: Actino (unclassified Actinobacteria), Chl-a (chlorophyll *a*), CyP (cyanobacterial plankton), DOC (dissolved organic carbon), EuAPP (eukaryotic autrotrophic picoplankton, i.e. pico-sized green algae), *Euglena* sp. (chlp5), ISS (inorganic suspended solids), POC (particulate organic carbon), Prok. cell number (prokaryotic cell number); for sample codes, see Supplementary Table S1.

### Changes in the prokaryotic community composition

In April 2013, Betaproteobacteria, especially the genus *Hydrogenophaga*, was the most abundant bacterial community member in the turbid pan (Fig. 1). Other groups with a high relative abundance (≥ 5%) were *Luteolibacter*, an uncultivated Planctomycetaceae genus, bacII and the acIII-A1 actinobacterial lineage. After continuous increase, the relative abundance of planktonic actinobacteria peaked in July (51%), with acIV, acTH1, acIII-A1 and *Nitriliruptor* as the most significant groups. Besides Actinobacteria, genus *Blastocatella* (Acidobacteria) was also abundant. In August and September 2013, in accordance with the microscopic observations, picocyanobacteria became abundant within the bacterial community, while the relative abundance of actinobacteria slowly decreased. From September 2013, the relative abundance of bacterial groups dominant during the winter/spring phytoplankton bloom started to increase. These taxa (i.e. *Rhodobaca* and another unidentified member of the alfVI lineage, *Luteolibacter*, various unidentified lineages from Cyclomorphaceae, Cryomorphaceae, Flavobacteriaceae and Saprospiraceae, genera *Flavobacterium* and *Aquiflexum* from Bacteroidetes, cyanobacteria, *Ilumatobacter*, acIII-A1, acSTL, and an unidentified group of Actinobacteria) with varying relative abundances dominated the planktonic microbial community until May 2014. From May, the relative abundance of planktonic actinobacteria increased suddenly, peaked in June with 89% within the bacterial community, and then started to decrease again by July. The acIII-A1 lineage was the most dominant group (54% of total bacteria) in June besides *Ilumatobacter*, acTH1, acSTL and *Nitriliruptor*.

The colored pan was dominated by Flavobacteria and Betaproteobacteria in the spring of 2013 (Fig. 1). In April, the genus *Flavobacterium* was the most plenteous (69%) in the water, and in the following months, other Bacteroidetes taxa (*Fluviicola*, *Algoriphagus*, unidentified Sphingobacteriales) were also abundant. From Betaproteobacteria, the genera *Hydrogenophaga* and *Limnohabitans* had high relative abundance. By the summer, Actinobacteria, especially the acIII-A1 lineage became the most significant constituent in the bacterial community. After refilling, an unidentified Cyclobacteriaceae group was the most abundant, with other taxa dominant during the winter/spring phytoplankton bloom (*Indibacter*, *Flavobacterium*, *Rhodobaca*, and other bacII and alfVI group representatives, and members of Verrucomicrobia and Actinobacteria). In December 2013 and January 2014, an unidentified Oligoflexaceae group was also abundant. *Hydrogenophaga* became abundant again within the community during the early months of 2014. Between April and June 2014, the relative abundance of the acIII-A1 actinobacterial lineage peaked again. In addition to this lineage, *Nitriliruptor*, an unidentified Actinobacteria, a Cryomorphaceae group, *Flavobacterium*, a Sphingobacteriales group, alf-B, alfVI and Comamonadaceae lineages were present with a high relative abundance.

### Effects of algae and zooplankton abundance on the planktonic bacterial community

In both pans, during the winter/spring algal blooms, members of the Cytophagia, Flavobacteriia and Rhodobacterales taxa dominated in the planktonic bacterial community. Later, in parallel with the increasing zooplankton abundance, algal blooms collapsed, and actinobacterial relative abundance peaks occurred (Fig. 3). Based on these observations we highlighted two groups within the microbial community: the algae-associated Cytophaga-Flavobacteriia-Rhodobacterales (CFR) group and planktonic actinobacteria. Changes in the relative abundance of the CFR group and actinobacteria in parallel with algal biomass and zooplankton abundance are presented in Fig. 3. The CFR group had a high relative abundance in the periods with high algal biomass, and further during the spring of 2013, when *Flavobacterium* was dominant within the community. The relative abundance of actinobacteria peaked during and after the peaks of microcrustacean abundance. It was especially striking in the turbid pan in May 2014, when both cladoceran and copepod microcrustaceans had an exceptionally high abundance with more than 10^4^ individuum per liter, and the relative abundance of planktonic actinobacteria (60%) also became surprisingly high. After the ‘microcrustacean peaks’, the abundance of rotifers increased in parallel with decreasing actinobacterial relative abundance.

**Fig. 3.**
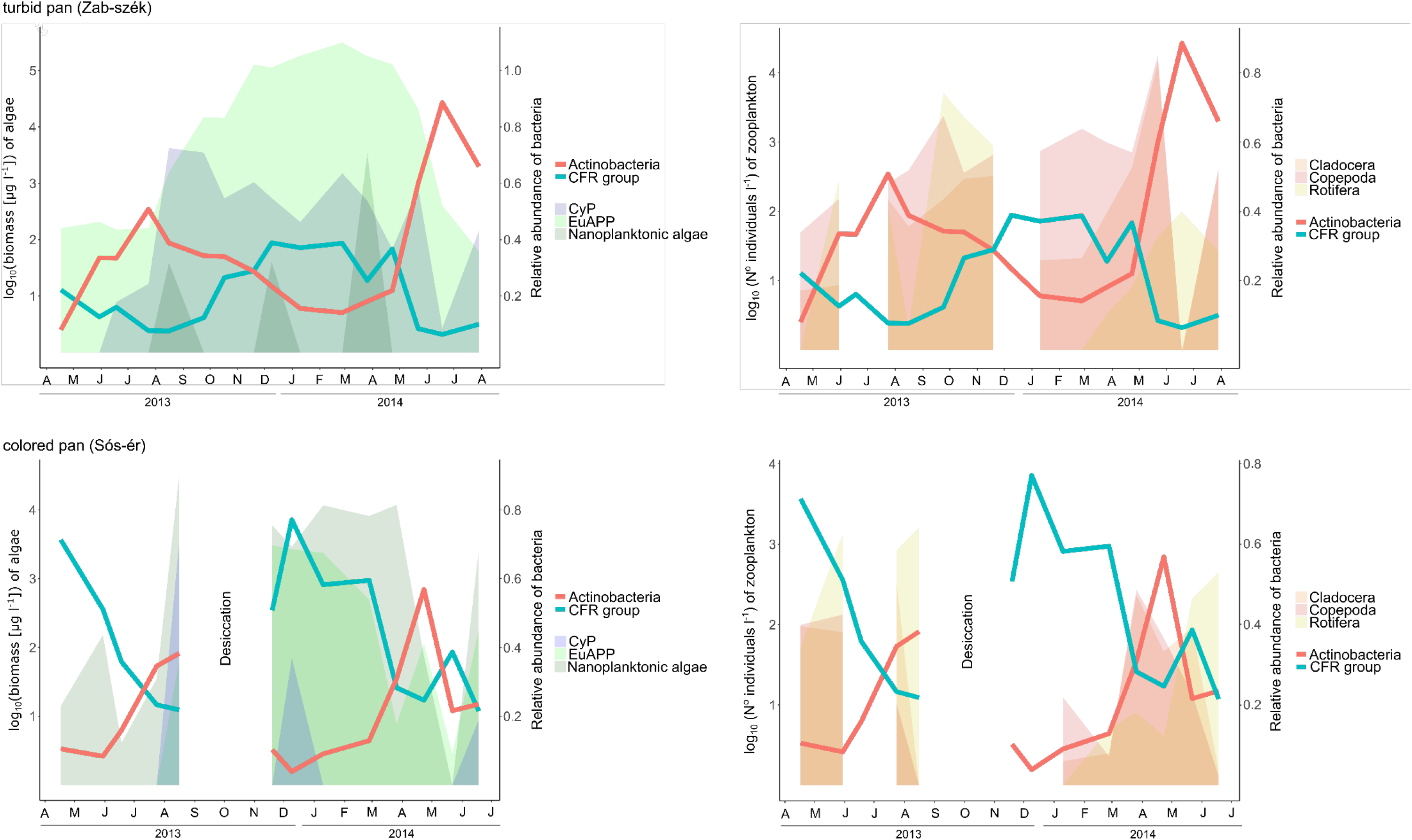
Relative abundance changes of Actinobacteria and the Cytophaga-Flavobacteria-Rhodobacterales (CFR) group, comparing to the biomass of planktonic algae and zooplankton abundance in the turbid and colored pan between April 2013 and July 2014. For visualization, algal biomass and zooplankton abundance values were transformed to a log10-scale. Letters on the x-axis are the abbreviations for months. Abbreviations: CyP (cyanobacterial plankton), EuAPP (eukaryotic autrotrophic picoplankton).

### Co-occurrence network analysis

To reveal causal relations and correlations within the microbial community of the pans and between the bacterioplankton and environmental factors, co-occurrence networks were created based on time-shifted and local correlations. This resulted in a single interconnected network in the case of both pans (Fig. 4).

**Fig. 4.**
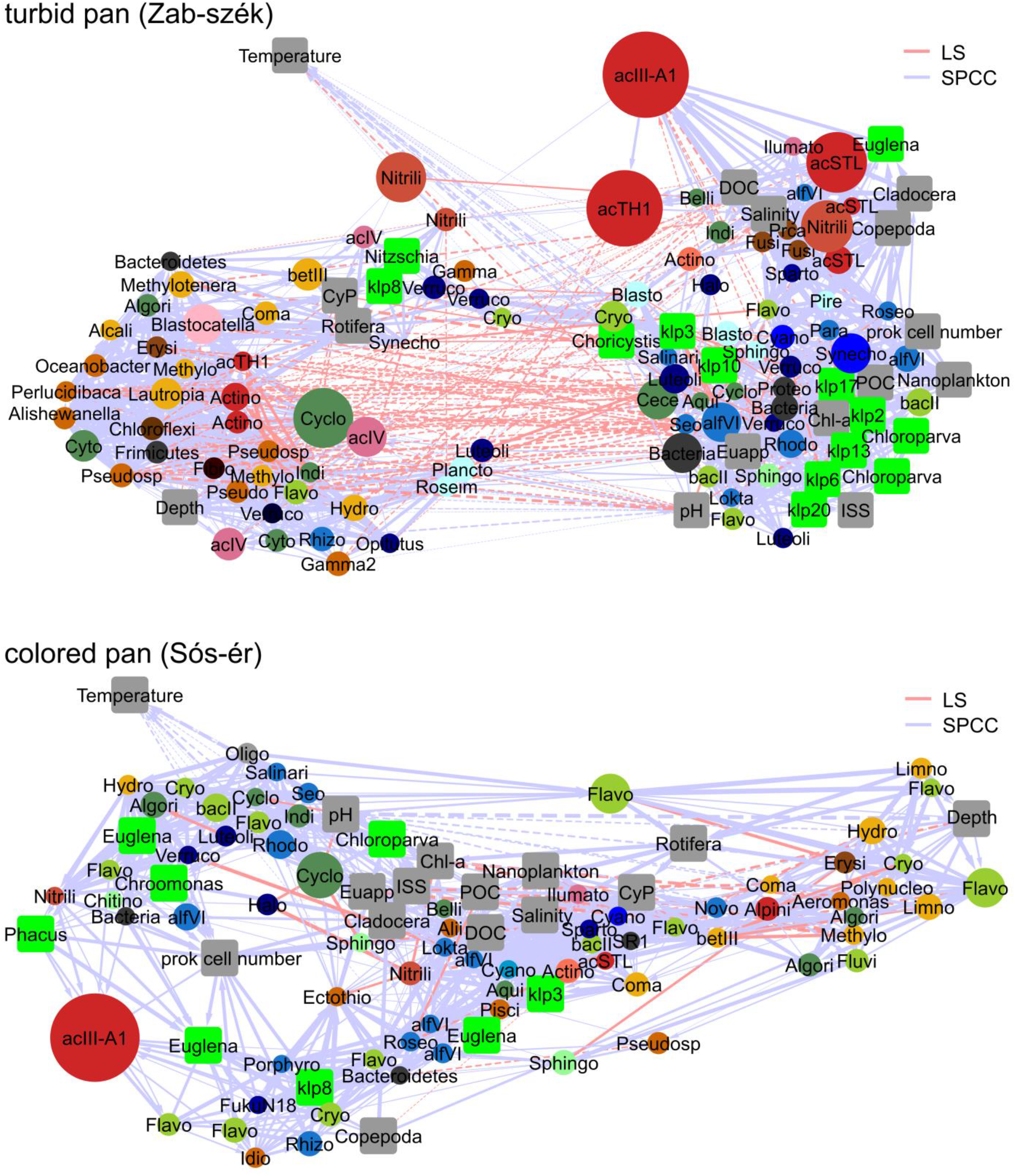
Co-occurrence networks of bacterial OTUs and environmental variables in the turbid and colored pans based on significant (p < 0.01, q < 0.01) time-shifted (blue edges) and local (red edges) correlations. Directed effects are shown with arrows, continuous lines represent positive correlations, and dashed lines depict negative correlations. Environmental parameters are shown in gray, identified algae chloroplast OTUs are shown with green squares. Names of the other hubs represent the closest assigned taxa to bacterial OTUs, and the size of the nodes correlates with the absolute abundance of the OTU in the sample set. The color of the nodes represents class-level phylogenetic relationships. Abbreviations: Actino (Actinobacteria), Alii (*Aliidiomarina*), Algori (*Algoriphagus*), Alpini (*Alpinimonas*), Aqui (*Aquiflexum*), Belli (*Belliella*), Blasto (*Blastopirellula*), Cece (*Cecembia*), Chl-a (chlorophyll *a*), Coma (Comamonadaceae), Cryo (Cryomorphaceae), Cyano (Cyanobacteria), CyP (cyanobacterial plankton), Cyclo (Cyclobacteriaceae), DOC (dissolved organic carbon), Ecto (*Ectothiorhodospira*), EuAPP (eukaryotic autrotrophic picoplankton), Flavo (*Flavobacterium*), Fibro (Fibrobacteraceae), Fluvi (*Fluviicola*), Gamma (TX1A55 Acidithiobacillales), Gamma2 (K189A clade Gammaproteobacteria), Halo (*Haloferula*), Hydro (*Hydrogenophaga*), Idio (*Idiomarina*), Illumato (*Ilumatobacter*), Indi (*Indibacter*), ISS (inorganic suspended solids), Limno (*Limnohabitans*), Lokta (*Loktanella*), Luteoli (*Luteolibacter*), Nitrili (*Nitriliruptor*), Novo (*Novosphingobium*), Oligo (Oligoflexaceae), Para (*Paracoccus*), Pisci (Piscirickettsiaceae), POC (particulate organic carbon), Porphyro (*Porphyrobacter*), Prca (*Proteocatella*), Prok. cell number (prokaryotic cell number), Pseudo (*Pseudomonas*), Pseudosp (*Pseudospirillum*), Roseim (*Roseimaritima*), Roseo (*Roseococcus*), Rhodo (*Rhodobaca*), Seo (*Seohaeicola*), Sparto (Spartobacteria), Synecho (*Synechococcus*), Verruco (Verrucomicrobia).

In the turbid pan, from the 7,260 correlations tested, 385 (LS) and 1,246 (SPCC) were significant. The number of shared edges among these correlations was 237. The resulting network contained 120 nodes. The density of the network was 0.2, and the centralization was 0.152. The following nodes had the highest number of edges: chlorophyll *a* (41), OTU50 (Actinobacteria) (39), OTU7 (*Rhodobaca*) (38) and OTU78 (TK10-Chloroflexi) (38). The average number of neighbors of the nodes was 23.2. Chlorophyll *a* (0.045), OTU7 (*Rhodobaca*) (0.039), OTU1 (acIII-A1) (0.031), OTU34 (*Seohaeicola*) (0.030), OTU11 (acSTL) (0.026) and OTU26 (*Synechococcus*) (0.026) had the highest betweenness centrality values.

Within the colored pan samples, from the 4,656 correlations tested, 97 (LS) and 859 (SPCC) were significant, and 63 correlations were shared between the two approaches. The network contained 95 nodes, the network density was 0.2, with a centralization of 0.154. The highest significant correlations were detected with nodes OTU151 (*Aliidiomarina*) (33), OTU69 (*Loktanella*) (33), POC and salinity (31-31). Node’s average number of neighbors was 18.8. The highest betweenness centrality values were obtained with OTU69 (*Loktanella*) (0.029), OTU3 (*Nitriliruptor*) (0.021) and OTU10 (*Flavobacterium*) (0.016).

On the resulting edge-weighted arrangement of the networks, the temperature was separated from other nodes. On the co-occurrence network of the turbid pan, two major group of nodes were separated from each other, with many negative correlations: (1) OTUs presented during higher water levels, mostly from the spring of 2013 (these were, for example, beta- and gammaproteobacteria, freshwater actinobacteria, *Blastocatella*), and (2) an algae-associated microbial group containing picoeukaryotic algae and cyanobacteria, the previously mentioned CFR-group and Verrucomicrobia. With the second group, OTU1 (acIII-A1) was associated with a number of directed correlations; chloroplast and CFR OTUs effected OTU1 negatively. On the other hand, cladoceran (SPCC: 0.92) and copepod abundance (SPCC: 0.89), and from the more abundant OTUs, OTU17 (*Nitriliruptor*) (SPCC: 0.91), and OTU45 (Spartobacteria) (SPCC: 0.95) had positive effects. OTU1 had a strong positive correlation with chloroplast OTU5 (*Euglena* sp.) (SPCC: 0.90), DOC (SPCC: 0.86), salinity (SPCC: 0.76), OTU4 (*Ilumatobacter*) (SPCC: 0.82) and OTU11 (acSTL) (SPCC: 0.67), and had a positive effect on OTU5 (acTH1) (SPCC: 0.85) and OTU3 (*Nitriliruptor*) (SPCC: 0.72). OTU11 and OTU81 (both acSTL) had the same relationship with the environmental parameters, while OTU51 (acSTL) had a positive effect on OTU5 (acTH1) and was associated with cladocerans and copepods, indicating that it was abundant prior to the other two acSTL OTUs. Between the two major groups of the turbid pan network, OTU2 (*Synechococcus*), chloroplast OTU8 (Bacillariophyta) and OTU9 (*Nitzschia*) were situated and had a positive effect on the rotifers. These groups dominated in the water during the summer period. The *Nitriliruptor* OTU3 also had a positive effect on these groups.

The colored pan co-occurrence network contained fewer negative associations, and it was more integrated than the turbid one. Most of the environmental parameters were situated in a central position. The CFR group and certain chloroplast OTUs were associated with each other, while the dominant OTUs during the high water level (mostly flavobacteria and betaproteobacteria) were separated loosely from other groups. OTUs belonging to the same genus were associated with different environmental factors in some cases, for example OTU9 (*Flavobacterium*) was associated with other bacterial groups abundant in the samples collected at high water levels in spring, while OTU71 (*Flavobacterium*) was connected with algae-associated groups, and OTU10 (*Flavobacterium*) was associated with both groups. In the colored pan, copepods (SPCC: 0.78), chloroplast OTU15 (*Phacus* sp.) (SPCC 0.82) and chloroplast OTU8 (Bacillariophyta) (SPCC: 0.83) had a positive effect on OTU1 (acIII-A1), which was also connected with chloroplast OTU18 (*Euglena* sp.) (SPCC: 0.82) and the prokaryotic cell number (SPCC: 0.80). Rotifers in the colored pan had an effect on the phytoplankton, while Cladocera and Copepoda had a strong positive correlation with each other, both in the turbid (SPCC: 0.99) and in the colored pan (SPCC: 0.93).

## Discussion

The environmental variables measured reflected multiple extreme conditions and were similar to those that had previously been reported in soda pans of the Carpathian Basin (Boros *et al.*, 2014, 2017); i.e. the turbid Zab-szék pan had a higher average concentration of ISS (1436 mg l^−1^) than the colored Sós-ér pan (86 mg l^−1^), while DOC values where higher in the latter (75 mg l^−1^ vs. 490 mg l^−1^, respectively), pH was alkaline (~9-10) and the salinity ranged between 1.5 to 18.9 g l^−1^ (subsaline to hyposaline, according to Hammer, 1986). Prokaryotic cell numbers were in the same magnitude as had been measured previously in the soda pans of the region studied (10^6^-10^8^ cells ml^−1^) (Vörös *et al.*, 2008), but higher than in five soda pans from the Seewinkel region (Austria) (10^6^ cells ml^−1^) (Eiler *et al.*, 2003). Interestingly, the number of cells did not increase linearly with depth, proving the dynamic nature of the prokaryotic community and the ecosystem. Previous studies have shown that picoeukaryotic algae have a competitive advantage at low temperature and low light intensity values, and, therefore, winter blooms of these small algae have frequently been observed in soda pans before (Somogyi *et al.*, 2009; Pálffy *et al.*, 2014). Despite the fact that the pans studied represented different ecotypes (of ‘turbid’ and ‘colored’ character), the same prokaryotic groups were present in both pans. Moreover, similar seasonal patterns were observed in the microbial community structure. Certain groups of planktonic actinobacteria were characteristic in both of the pans studied: the acIII-A1, acSTL, acTH1 groups and genera *Nitriliruptor* and *Ilumatobacter*. The classes Cytophagia, Flavobacteria and the order Rhodobacterales (CFR group) were also abundant, mostly during algal blooms. Betaproteobacteria, which is a well-known and abundant group in various freshwater ecosystems (Newton *et al.*, 2011), were frequent at higher water levels and lower salinity. Picocyanobacteria (represented by the genus *Synechococcus*) occurred frequently in the lakes, which also corresponded with literature data and with previous studies on the soda pans within this region (Felföldi *et al.*, 2009, 2011; Somogyi *et al.*, 2009); however, sometimes filamentous groups (e.g., *Anabaenopsis*) were also detected. Several uncultured/unclassified taxa were abundant in the pans and were important constituents of the community.

Compared to other alkaline lakes (e.g., Vavourakis *et al.*, 2016), the presence of Archaea was not significant (13%) within the prokaryotic community.

Planktonic actinobacteria were one of the most significant bacterial groups in the pans studied, reaching even 89% relative abundance within the bacterial community, which was the highest value compared to any literature data from aquatic habitats. Especially abundant (up to 54%) was the currently poorly characterized acIII-A1 lineage, which was first described based on clone sequences from the hypersaline Mono Lake soda lake (California, USA) and the meromictic Lake Sælenvannet estuary (Norway) (Warnecke *et al.*, 2004). Later, Ghai and colleagues (2012) also identified this lineage in coastal lagoons. We suspect that the main reason behind the lack of reports on this group is that they could be only distinguished by the FreshTrain dataset, although other methodological problems (e.g., the widely used 1492R primer fails to amplify some groups of Actinobacteria; Farris and Olson, 2007) could also have contributed to this phenomenon. Using ARB-SILVA, the taxonomic assignment for these reads results in an ‘unclassified’ rank within the family Microbacteriaceae. To test our hypothesis, we reanalyzed some previously published datasets from the epilimnion of saline-alkaline lakes (SRA: SRS950620 – Baatar *et al.*, 2016, SRA: SRP044627 –454 FLX Titanium runs, Sinclair *et al.*, 2015, SRA: SRR10674068, Szabó A, Márton Zs, Boros E, Felföldi T, unpublished results) downloaded from the NCBI SRA, applying the same sequence analysis pipeline as for our samples (data not shown). As a result, the acIII-A1 group was detected with high relative abundance in three soda lakes in the Seewinkel region, Austria (Silber Lacke: 21%, Unterer Stinker: 6%, Zicklacke: 7%), from Lake Teniz in North-Kazakhstan (11%) and from the surface water of Lake Oigon (5%), which is another saline steppe lake with high pH in Mongolia.

Based on our findings, the high grazing pressure of microcrustaceans coincided with the unusually high relative abundance of acIII-A1 and other planktonic actinobacterial groups during the spring/summer transition in both pans. Due to their small cell size (< 0.1 μm^3^) and the presence of S-layer in their cell wall, planktonic actinobacteria are more resistant to grazing compared to other freshwater bacteria (Tarao *et al.*, 2009; Newton *et al.*, 2011). Pernthaler *et al*. (2001) proved experimentally that due to the presence of a size-selective predator (*Ochromonas* nanoflagellate), the relative abundance of the acI planktonic actinobacterial lineage, which are among the most abundant bacteria in temperate lakes (Newton *et al.*, 2011), increased suddenly and even reached 60% in the model freshwater community. Hahn *et al*. (2003) obtained similar results with the Luna actinobacterial clades. In another study, Šimek *et al*. (2018) compared the population growth of heterotrophic nanoflagellates (HNF) on different freshwater betaproteobacterial and acIII-A1-related Luna2 actinobacterial strains. According to their findings, a rapid HNF population growth occurred on all bacterial strains tested, except for the Luna2 actinobacterium. Tarao and colleagues (2009) proved that the destruction of the actinobacterial cell wall’s surface layer results increased predation on Luna2 strains. In certain circumstances, especially following a phytoplankton spring bloom, metazoan grazing could have a greater impact on bacterial diversity than the predation by HNFs (Hahn and Höfle, 2001; Langenheder and Jürgens, 2001). Studies have shown that microcrustaceans and rotifers also like to feed on bacterioplankton, not just on algae (Sanders *et al.*, 1989; Turner and Tester, 1992; Roff *et al.*, 1995; Kirchman, 2002;, Agasild and Nõges, 2005; Berga *et al.*, 2015). *Daphnia magna*, an abundant microcrustacean in these soda pans (Tóth *et al.*, 2014), usually feeds on larger bacterial cells but also consumes HNFs, ciliates and algae (Jürgens, 1994; Berga *et al.*, 2015). During an experiment using aquatic metacommunities (Berga *et al.*, 2015), in the presence of *Daphnia magna*, the relative abundance of Actinobacteria and Sphingobacteria increased. In our study, during the period investigated, *Daphnia* was also present in the pans, but the most abundant cladoceran species (especially during late spring and early summer) was *Moina brachiata*, which is a characteristic species of soda pans of the Carpathian Basin (Tóth *et al.*, 2014). We did not find any data in the literature on the relationship of *Moina brachiata* and bacterioplankton; however, a closely-related cladoceran, *Moina macrocopa*, was the most efficient predator of bacteria in an experiment where *Vibrio cholerae* cells were used (Ramirez *et al.*, 2012). The other abundant microcrustacean species was *Arctodiaptomus spinosus*, a characteristic copepod in soda lakes (Tóth *et al.*, 2014). It mostly feeds on algae, protists and rotifers; however, in the nauplius stage, copepods could also be bacterivorous (Turner and Tester, 1992; Roff *et al.*, 1995). It can be hypothesized that due to the large number of copepods, the abundance of the non-size selective bacterivorous ciliates (if they are present at all in remarkable number in the plankton) also decreases, thus favoring the growth of planktonic actinobacteria. Members of the acI clade, another small cell-sized planktonic actinobacteria, are capable of utilizing N-acetylglucosamine (monomeric unit of chitin and component of the bacterial murein cell-wall) (Beier and Bertilsson, 2011; Eckert et al., 2013; Garcia *et al.*, 2013), which can also contribute to the increase of their relative abundance with zooplankton numbers. In previous works on soda pans of the Seewinkel region (Eiler *et al.*, 2003; Krammer *et al.*, 2008), the authors stated that the observed high abundance of cladocerans (2 × 10^3^ specimen l^−1^) and protists must be an important regulatory factor on the bacterial population. In conclusion, intensive grazing by zooplankton could favor planktonic actinobacteria by the elimination of their larger, fast-growing bacterial competitors and also by their ability to degrade chitin, which could present in high amounts in the water by the end of the zooplankton ‘bloom’.

During or after the actinobacterial relative abundance peaks, the abundance of rotifers also increased in our study; however significant relationship between these groups was not proved statistically. Besides the small cell size, increasing the size is also a good strategy for a microbe to avoid predation (Hahn and Höfle, 2001). Probably this also contributes to the summer blooms of filamentous cyanobacteria in these pans (Boros *et al.*, 2013).

In addition to actinobacteria, the other abundant taxon was the Cytophaga-Flavobacteria group in the pans studied. In aquatic environments, the CF group usually participates in the degradation of dissolved and particulate organic matter; it is capable of utilizing biopolymers that have a high molecular weight, which are frequently associated with algal blooms (Kirchman, 2002; Newton *et al.*, 2011; Teeling *et al.*, 2012; Williams *et al.*, 2013; Buchan *et al.*, 2014). Flavobacteria were also abundant in the colored pan at higher water levels. These microorganisms are not just capable of utilizing algae-derived compounds, but also macrophyte-derived organic matter, which could originate from the allochthonous terrestrial and autochthonous marshland vegetation of the pan (Boros et al, 2020). In a preliminary metagenomic study (Szabó *et al.*, 2017), genes of the TonB receptor system were the most abundant genes within the planktonic community, which highlights the importance of the CF group and algae-derived organic compounds in nutrient cycling. Members of Rhodobacterales (alfVI) were also abundant during the winter-spring algal bloom in the pans. These microbes are related to phytoplankton-derived organic matter but rather utilize algal exudates instead of larger compounds (Teeling *et al.*, 2012; Williams *et al.*, 2013; Buchan *et al.*, 2014). In the co-occurrence network of the turbid pan, the chlorophyll *a* content had the highest number of connections and betweenness centrality values (the second highest was a Rhodobaca-related OTU, which was also correlated with algal biomass). These correlations were distorted in the colored pan, because of the contribution of macrophyton-derived organic matter to the total carbon budget.

During the spring of 2013, the high water level reduced the pH and salinity of the pans. At this stage, the colored pan rather resembled an alkalic ‘freshwater’ marsh, and this was also reflected in the composition of the bacterial community. During this phase, mostly alkalitolerant heterotrophic groups were abundant (Flavobacteriia, *Novosphingobium*), instead of the aforementioned typical soda lake bacterial groups, which were otherwise present in the pans. The former taxa are capable of utilizing plant-derived matter, and they had high growth rates in natural aquatic habitats. Therefore, they are able to become dominant quickly in the community, decreasing the overall bacterial diversity. Thus, besides zooplankton grazing and algal blooms, the water depth also has a remarkable impact on the structure of the planktonic microbial community in the soda pans studied.

In conclusion, even if the prokaryotic communities of the colored Sós-ér and the turbid Zab-szék pans differed significantly, the same core-taxa were dominant in both pans. These groups are characteristic for other saline-alkaline environments worldwide, especially for soda lakes; however, the importance of acIII-A1 actinobacteria in the bacterioplankton of lakes was revealed for the first time in this study. Seasonal abiotic and biotic effects (predation, algal blooms and water depth) have a great impact on the microbial composition, and our study highlights the dynamic nature of the prokaryotic community in soda lakes and the importance of frequent sampling to reveal ecological processes. Since most soda lakes are situated in remote places of the world, soda pans of the Carpathian Basin are ideal sites to study these environments in detail. Furthermore, these moderately saline, hypertrophic habitats with rapidly changing environmental conditions and a simple food web structure may be regarded as model systems of the current and predicted processes associated with global climate change and the growing human population (introduction of salt water to freshwater, weather changes, a loss of biological diversity and pollution of inland waters). Moreover, further studies are needed to reveal the metabolic potential and geographic distribution of acIII-A1 actinobacteria, the composition of protists and the controlling role of viruses on the planktonic microbial community in soda lakes and pans.

## Experimental procedures

### Sample collection

A detailed description of the pans studied (the turbid Zab-szék pan and the colored Sós-ér pan; Supplementary Fig. S1) can be found in Boros *et al.* (2016) and in Szabó *et al.* (2017). Samples were collected monthly from 17 April 2013 to 29 July 2014. The Sós-ér pan was completely desiccated on 24 September, 17 October 2013 and after 18 June 2014, and, therefore sampling on these dates was not possible. Based on our investigation on spatial heterogeneity, using three replicate samples from these sites, there were no remarkable differences in the bacterial community composition (Bedics *et al.*, 2019), presumably due to the constant mixture of the shallow water by the wind (Boros *et al.*, 2017). Nevertheless, to represent the total bacterial community of the pans, composite samples were taken from at least ten different points each time in the vicinity of the deepest parts of the open water bodies. To avoid diel differences in community composition and environmental parameters as referred to by Kirschner and colleagues (2002), samplings were performed approximately the same time on every occasion. Further information about sample collection dates is given in Supplementary Table S1. The determination of physicochemical parameters, prokaryotic cell count, composition and biomass of algae was performed as described in Mentes *et al.* (2018). Salinity values were calculated from electric conductivity based on the equation published by Boros *et al.* (2014). To determine mesozooplankton abundance, a 10 l water sample was randomly collected from the open water of each pan and filtered through a 30 μm mesh plankton net, then preserved in 70% ethanol (zooplankton was not enumerated in the case of samples collected on 18 June 2013 and on 9 Dec 2013 due to technical problems).

### Community DNA isolation, amplification, sequencing

DNA extraction, 16S rRNA gene amplification and amplicon sequencing were performed as described in Szabó *et al.* (2017), while qPCR measurements to determine the ratio of archaea to bacteria were carried out as described in Korponai *et al.* (2019).

### Bioinformatic and statistical analyses

Sequence processing, taxonomic assignments and OTU-picking (defined at a 97% similarity level) were carried out using mothur v1.35 (Schloss *et al.*, 2009), as described in Szabó *et al.* (2017) (scripts are also available in the Supplementary Material of this paper). For abundant unassigned OTUs, the FreshTrain reference dataset (18 Aug 2016 release, Newton *et al.*, 2011) was used as a taxonomic reference. Cyanobacterial and chloroplast sequences were also analyzed separately from bacterial reads; the OTU’s taxonomic identification was carried out based on phylogenetic analysis with 16S rRNA gene sequences obtained from GenBank and from reference strains (using the methods described in Felföldi *et al.* (2011) and Borsodi *et al.* (2017); see additional details in Supplementary Table S2).

Subsampling the reads to the smallest data set (n = 1896), calculation of richness (sobs - number of OTUs at a 97% similarity threshold, Chao1 - Chao 1 index and ACE - Abundance-based Coverage Estimator), and diversity estimators (Simpson’s inverse and Shannon indices), along with the Bray-Curtis similarity index, were also performed using mothur. For the determination of significant differences in community composition between pans and among seasons, a two-way PERMANOVA test was applied based on Bray-Curtis similarity with 9999 permutations using the PAST3 program (Hammer *et al.*, 2001). PAST3 was also used to perform a SIMPER test, which determined the OTUs responsible for the dissimilarity among all of the samples. Non-metric multidimensional scaling (NMDS) ordinations and ‘envfit’ analyses were carried out with R (R Core Team, 2017) using the vegan (Oksanen *et al.*, 2017) and ggplot2 (Wickham 2009) packages. All variables were log10-transformed before the ‘envfit’ analysis to reduce the effect of outliers and to obtain normal distribution of the data. Shapiro-Wilk tests were used to check normality.

Co-occurrence networks were created using the eLSA program (Xia *et al.*, 2011, 2013) to reveal local and time-shifted correlations among abundant (with at least 1% relative abundance in one sample) bacterial and chloroplast OTUs and environmental parameters within each pan. Data were normalized using the percentileZ function, and p-values were estimated based on 1000 permutations. Strongly significant (q < 0.01, p < 0.01) local similarity score (LS) and delay-shifted Pearson’s correlation coefficient (SPCC) parameters were used as edges of the networks. Cytoscape v3.4.0 was used for network visualization and analysis (Shannon *et al.*, 2003). To indicate relationships between nodes, networks were generated using the edge-weighted spring-embedded layout. Topological metrics were calculated using the network analysis plug-in (Assenov *et al.*, 2008).

## Data availability

Raw sequence data were submitted to NCBI under BioProject ID PRJNA272672.

## Acknowledgements

The authors are thankful to Balázs Németh and Tamás Sápi for their assistance during sampling, to Csaba Ferenc Vad, Vera Senánszky, Adrienn Tóth and Katalin Zsuga for zooplankton data, to Zsuzsanna Márton and Anikó Mentes for their help in the qPCR measurements and the sequencing of algal strains, respectively, and further, to Susann Pihl for proofreading the manuscript.

## Contributions

A.Sz. and T.F. wrote the paper. B.S. and E.B. carried out the sampling and determined physicochemical parameters. B.S. and L.V. determined algal cell counts and biomass. N.Sz-T. determined prokaryotic cell counts. Zs.H. supervised the determination of zooplankton abundance and identification of taxa. A.Sz. and K.K. performed DNA isolation, amplification and sequencing. A.Sz. carried out qPCR measurements, bioinformatic analyses and the statistical analyses with B.V. T.F and K.M. advised on the interpretation of the results. All authors read, commented and validated the final version of the manuscript.

---

**Supplementary Fig. S1.**
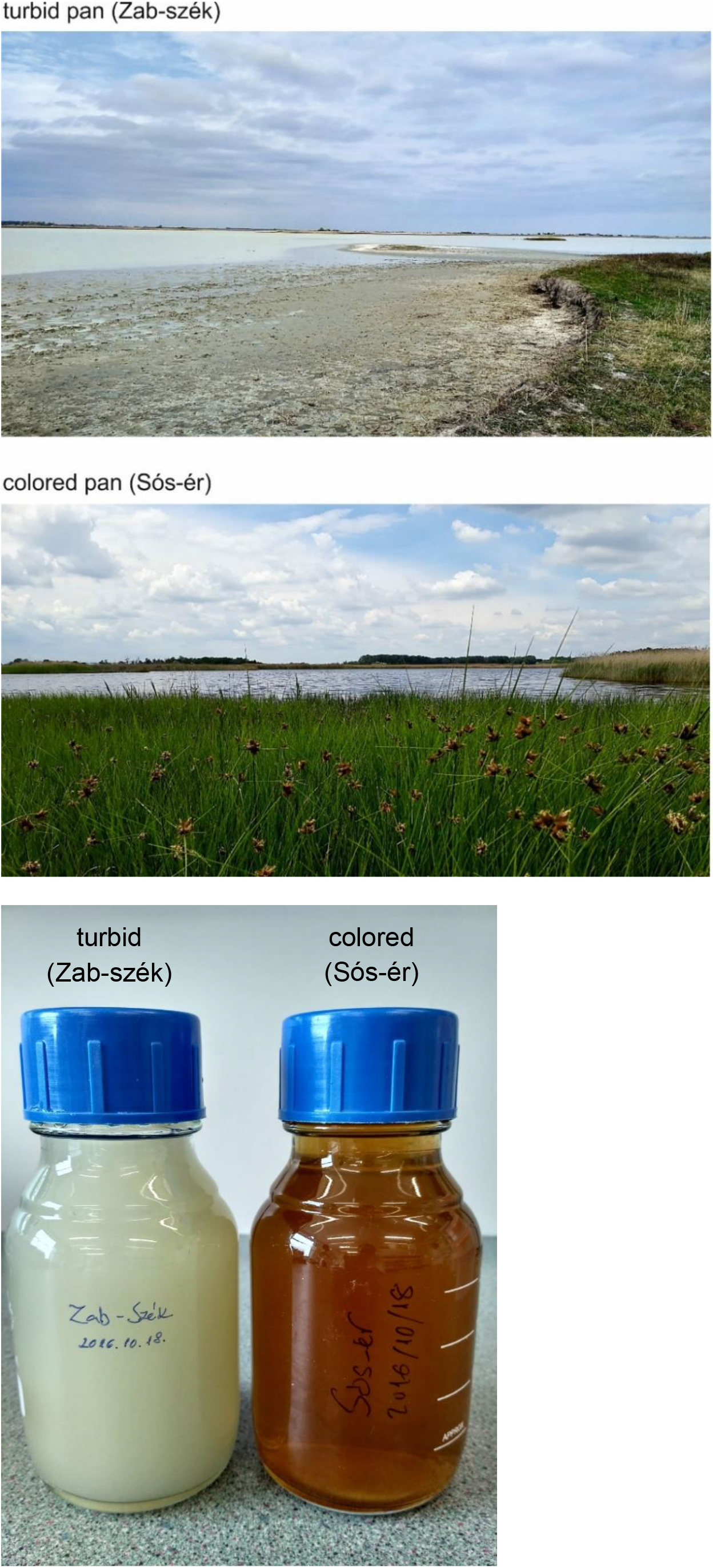
View of sampling sites and the color of their water.

**Supplementary Fig. S2.**
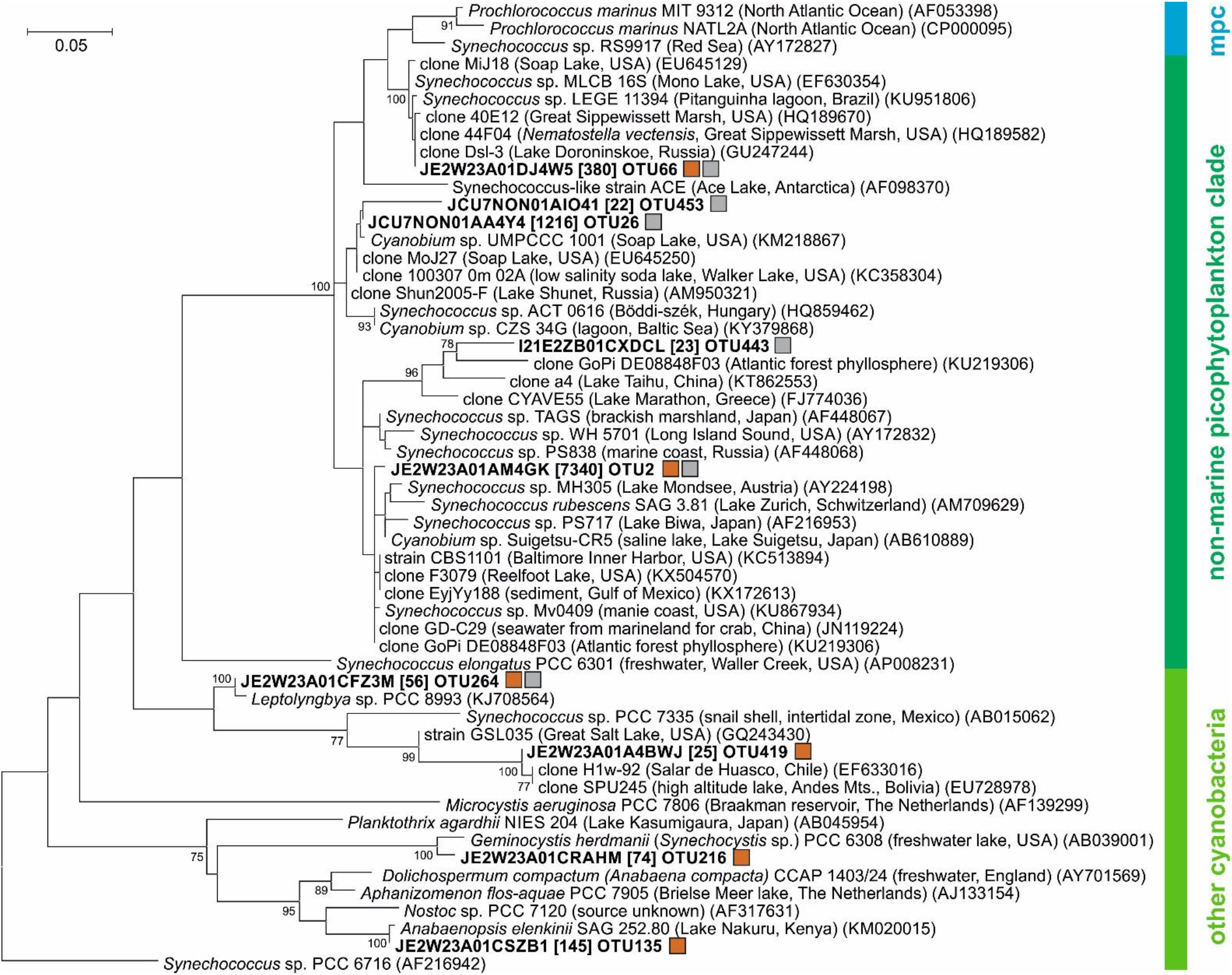
Phylogenetic position of cyanobacterial 16S rRNA gene amplicon reads from the soda pan samples collected between April 2013 and July 2014. The tree was constructed using the maximum likelihood method and is based on 365 nucleotide positions. Each OTU is represented with the most abundant read. Read numbers are given in brackets (only OTUs having at least 20 reads in the whole dataset are shown). Bootstrap values higher than 70 are shown at the nodes. OTUs detected in the colored and turbid pan are marked with brown and gray squares, respectively. mpc – marine picophytoplankton clade.

**Supplementary Fig. S3.**
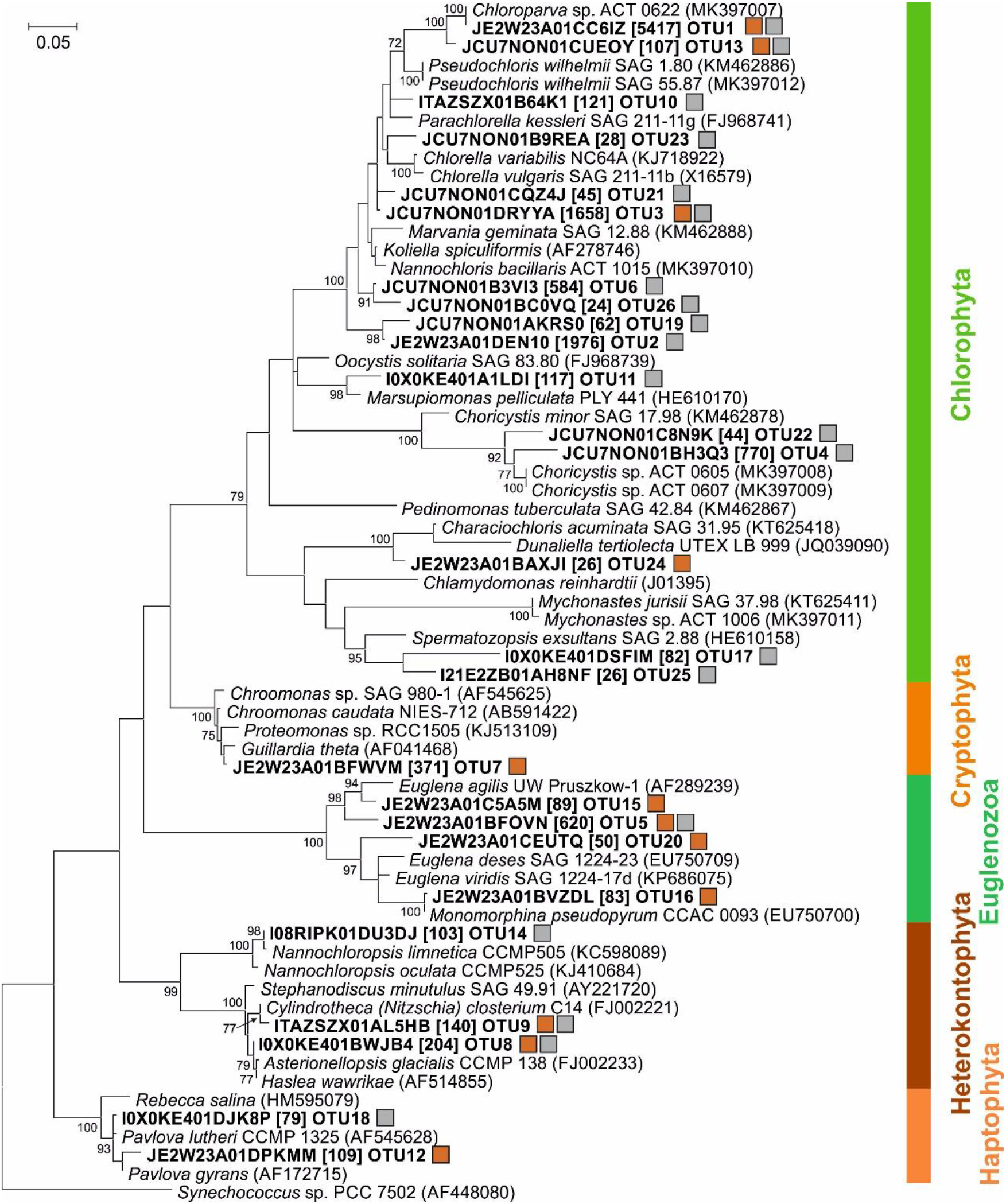
Phylogenetic position of chloroplast 16S rRNA gene amplicon reads from the soda pan samples collected between April 2013 and July 2014. The tree was constructed using the maximum likelihood method and is based on 439 nucleotide positions. Each OTU is represented with the most abundant read. Read numbers are given in brackets (only OTUs having at least 20 reads in the whole dataset are shown). Bootstrap values higher than 70 are shown at the nodes. OTUs detected in the colored and turbid pan are marked with brown and gray squares, respectively.

**Supplementary Fig. S4.**
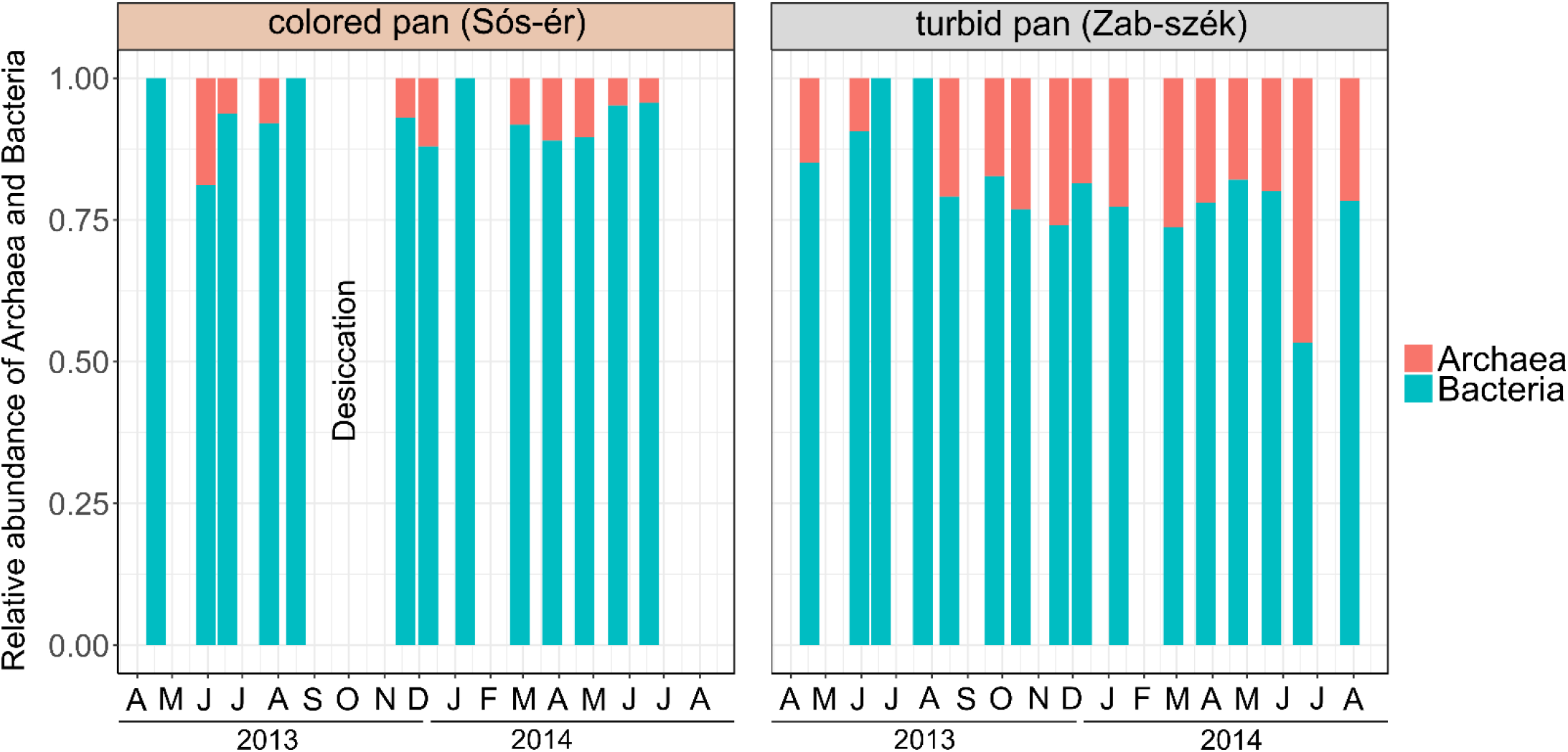
Relative abundance of Archaea and Bacteria in the colored pan (Sós-ér) and turbid pan (Zab-szék pan) between April 2013 and July 2014. Letters on the x-axis are the abbreviations for months.

**Supplementary Fig. S5.**
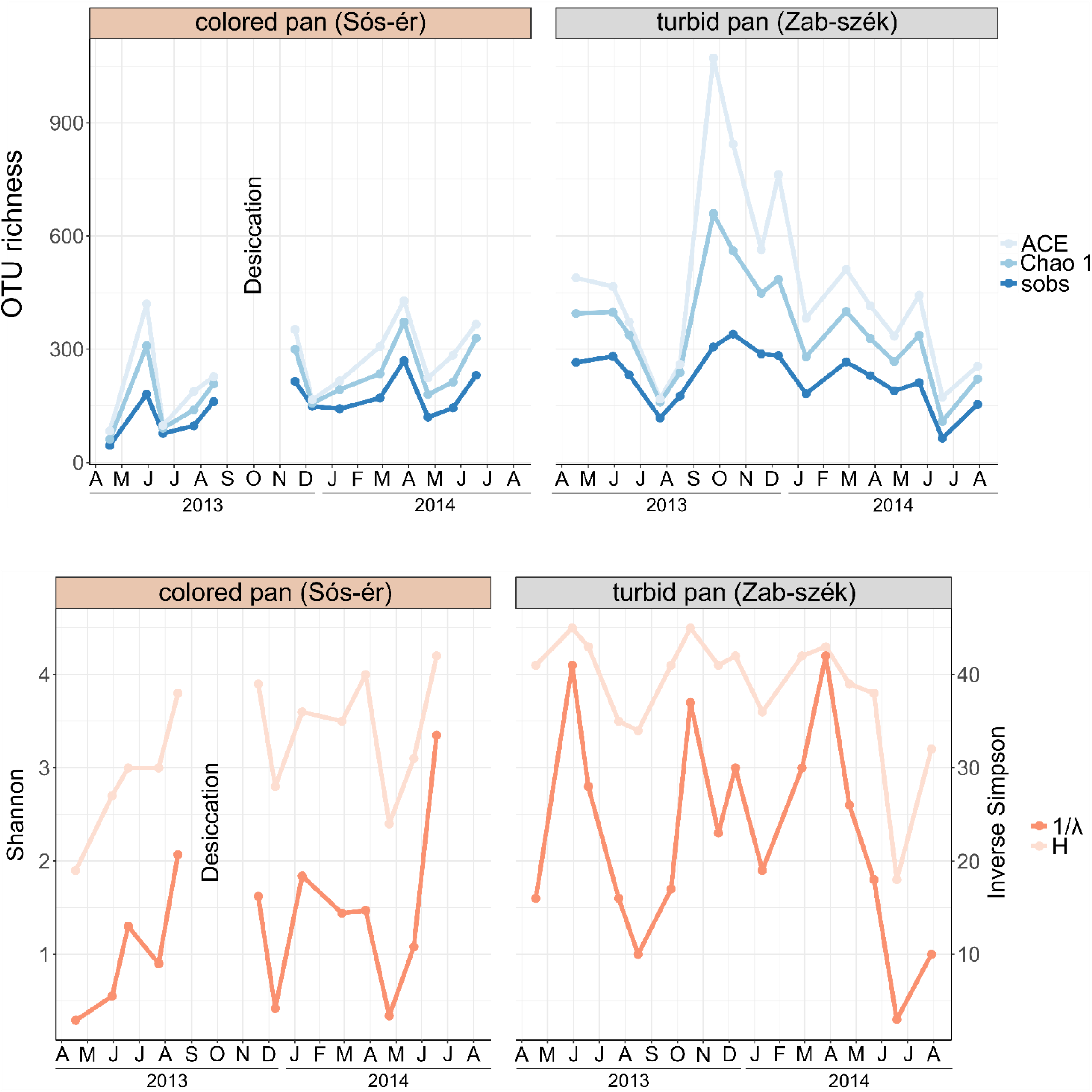
Changes in OTU richness and diversity in the studied pans between April 2013 and July 2014. Letters on the x-axis are the abbreviations for months. Abbreviations: H (Shannon diversity index), 1/λ (inverse Simpson diversity index).

**Supplementary Fig. S6.**
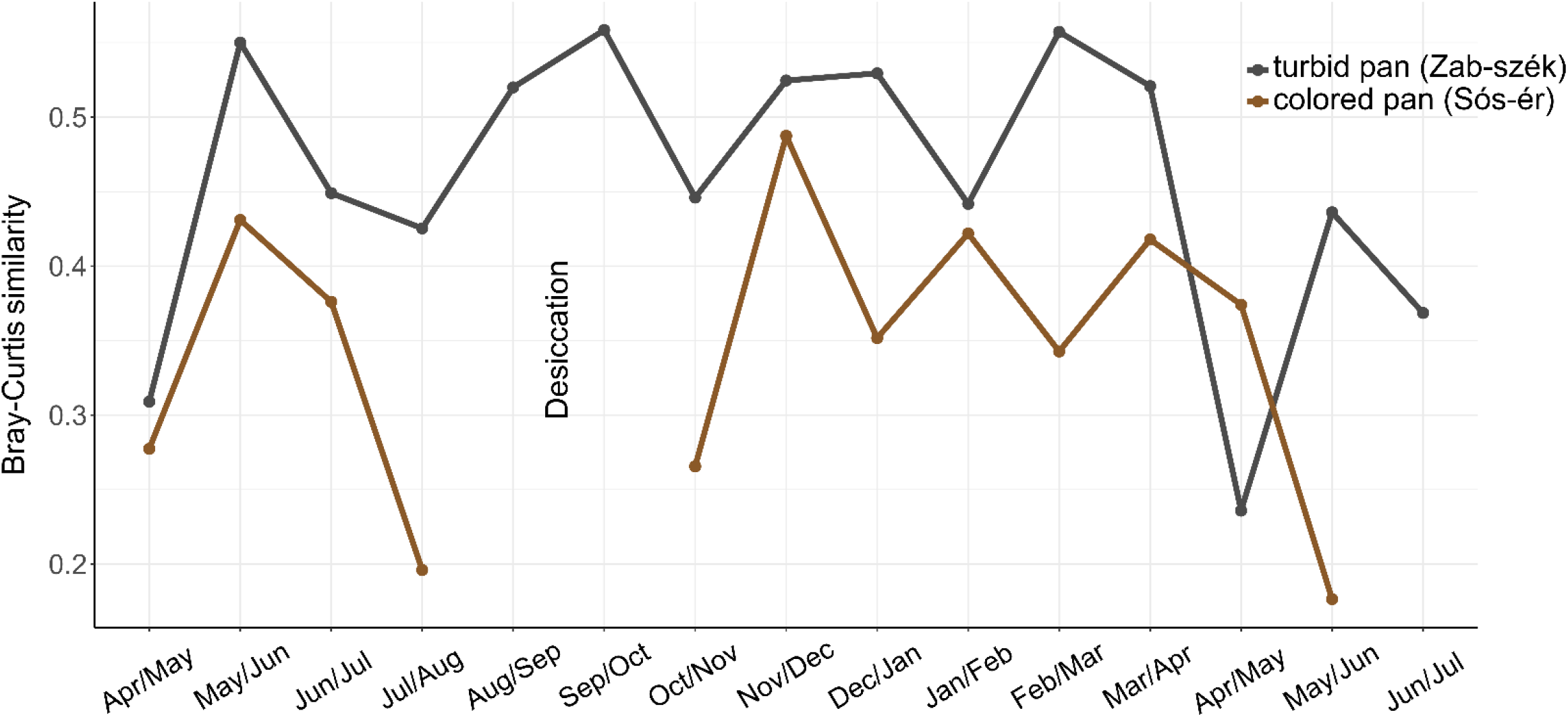
Monthly changes in Bray-Curtis similarity within bacterial communities of the pans studied

**Supplementary Table S1.** Physicochemical characteristics and selected biological variables of the water samples studied

| Sample code | Sampling date | Depth<br>cm | Temperature<br>°C | pH | Conductivity<br>μS cm<sup>-1</sup> | Chlorophyll *a*<br>μg l<sup>-1</sup> | Salinity<br>g l<sup>-1</sup> | DOC<br>mg l<sup>-1</sup> | POC<sup>1</sup><br>mg l<sup>-1</sup> | ISS<sup>1</sup><br>mg l<sup>-1</sup> | Prokaryotic<br>cell number<br>cells ml<sup>-1</sup> | CyP<sup>2</sup><br>biomass<br>μg l<sup>-1</sup> | EuAPP<sup>3</sup><br>biomass<br>μg l<sup>-1</sup> | Nanoplanktonic<br>algal biomass μg l<sup>-1</sup> | Rotatoria<br>individuum l<sup>-1</sup> | Cladocera<br>individuum l<sup>-1</sup> | Copepoda<br>individuum l<sup>-1</sup> |
| --- | --- | --- | --- | --- | --- | --- | --- | --- | --- | --- | --- | --- | --- | --- | --- | --- | --- |
| SO02 | 17 April 2013 | 50 | 17 | 8.3 | 1880 | 1.5 | 1.5 | 84 | 1 | 3 | 2,39E+07 | 0 | 0 | 14 | 52 | 95 | 99 |
| SO03 | 30 May 2013 | 45 | 17 | 8.5 | 2980 | 3.6 | 2.4 | 144 | 4 | 24 | 1,42E+07 | 0 | 0 | 152 | 1352 | 80 | 133 |
| SO04 | 18 June 2013 | 35 | 30 | 8.7 | 3800 | 0.5 | 3.0 | 174 | 0 | 6 | 1,76E+07 | 0 | 0 | 4 | n.d. | n.d. | n.d. |
| SO05 | 24 July 2013 | 18 | 25 | 9.1 | 7850 | 1.8 | 6.3 | 350 | 5 | 7 | 2,27E+08 | 0 | 0 | 35 | 837 | 339 | 10 |
| SO06 | 16 August 2013 | 10 | 27 | 9.6 | 23600 | 463.0 | 18.9 | 1164 | 59 | 232 | 1,03E+08 | 3515 | 64 | 32150 | 1603 | 0 | 1 |
| SO09 | 19 November 2013 | 4 | 8 | 10.0 | 11400 | 348.9 | 9.1 | 626 | 46 | 173 | 7,24E+07 | 0 | 3077 | 5960 | 0 | 0 | 5 |
| SO10 | 09 December 2013 | 9 | 4 | 9.9 | 7790 | 340.0 | 6.2 | 533 | 13 | 92 | 3,14E+07 | 70 | 2681 | 2918 | n.d. | n.d. | n.d. |
| SO11 | 10 January 2014 | 9 | 5 | 9.9 | 9500 | 379.0 | 7.6 | 595 | 19 | 221 | 4,67E+07 | 0 | 2376 | 11670 | 0 | 0 | 12 |
| SO12 | 26 February 2014 | 18 | 7 | 9.1 | 4500 | 148.5 | 3.6 | 276 | 18 | 88 | 3,13E+07 | 0 | 494 | 8160 | 0 | 2 | 2 |
| SO13 | 26 March 2014 | 13 | 11 | 9.2 | 6800 | 64.9 | 5.4 | 449 | 10 | 19 | 4,74E+08 | 0 | 8 | 11840 | 8 | 234 | 272 |
| SO14 | 23 April 2014 | 8 | 21 | 9.3 | 10700 | 49.4 | 8.6 | 682 | 15 | 139 | n.d. | 0 | 114 | 75 | 4 | 46 | 68 |
| SO15 | 22 May 2014 | 13 | 25 | 9.3 | 7700 | 2.2 | 6.2 | 466 | 10 | 0 | n.d. | 0 | 3 | 0 | 209 | 93 | 14 |
| SO16 | 18 June 2014 | 7 | 24 | 9.6 | 15000 | 92.0 | 12.0 | 830 | 25 | 115 | 1,30,E+08 | 9 | 186 | 2710 | 451 | 1 | 1 |
| ZA02 | 17 April 2013 | 45 | 24 | 9.1 | 2430 | 6.8 | 1.9 | 31 | 6 | 1499 | 1,81,E+07 | 0 | 159 | 0 | 0 | 7 | 50 |
| ZA03 | 30 May 2013 | 31 | 17 | 9.3 | 3020 | 11.7 | 2.4 | 33 | 5 | 791 | 3,87,E+07 | 0 | 207 | 0 | 283 | 8 | 152 |
| ZA04 | 18 June 2013 | 32 | 26 | 9.2 | 3300 | 1.7 | 2.6 | 32 | 8 | 649 | 2,11,E+07 | 8 | 152 | 0 | n.d. | n.d. | n.d. |
| ZA05 | 24 July 2013 | 20 | 31 | 9.5 | 5500 | 3.8 | 4.4 | 50 | 4 | 372 | 2,96,E+07 | 16 | 159 | 0 | 115 | 149 | 256 |
| ZA06 | 16 August 2013 | 13 | 30 | 9.8 | 9410 | 44.1 | 7.5 | 75 | 9 | 447 | 1,02,E+08 | 4216 | 1594 | 39 | 2 | 61 | 393 |
| ZA07 | 24 September 2013 | 13 | 16 | 10.0 | 12300 | 153.2 | 9.8 | 92 | 36 | 2228 | 1,60,E+08 | 3516 | 14934 | 0 | 5148 | 150 | 2394 |
| ZA08 | 17 October 2013 | 6 | 17 | 10.1 | 8000 | 259.8 | 6.4 | 60 | 41 | 2031 | 6,88,E+07 | 542 | 14455 | 0 | 2304 | 296 | 361 |
| ZA09 | 19 November 2013 | 4 | 11 | 10.2 | 9400 | 430.2 | 7.5 | 74 | 41 | 1679 | 1,08,E+08 | 1053 | 126661 | 0 | 882 | 324 | 666 |
| ZA10 | 09 December 2013 | 6 | 4 | 10.2 | 6400 | 455.7 | 5.1 | 59 | 37 | 1115 | 6,57,E+07 | 549 | 114084 | 40 | n.d. | n.d. | n.d. |
| ZA11 | 10 January 2014 | 4 | 6 | 10.2 | 8000 | 1043.7 | 6.4 | 64 | 74 | 456 | 1,70,E+08 | 206 | 183349 | 0 | 0 | 20 | 742 |
| ZA12 | 26 February 2014 | 12 | 10 | 9.6 | 4800 | 1499.4 | 3.8 | 41 | 86 | 4438 | 5,34,E+07 | 1511 | 313554 | 0 | 0 | 21 | 1563 |
| ZA13 | 26 March 2014 | 7 | 14 | 9.7 | 6400 | 2555.5 | 5.1 | 53 | 127 | 1809 | 3,84,E+08 | 494 | 180479 | 3450 | 3 | 96 | 988 |
| ZA14 | 23 April 2014 | 3 | 25 | 9.7 | 9000 | 822.0 | 7.2 | 74 | 147 | 1527 | n.d. | 69 | 129407 | 0 | 8 | 428 | 710 |
| ZA15 | 22 May 2014 | 3 | 28 | 9.4 | 8300 | 59.6 | 6.6 | 97 | 55 | 3215 | 1,92E+08 | 824 | 20726 | 0 | 43 | 14160 | 17760 |
| ZA16 | 18 June 2014 | 2 | 28 | 9.9 | 21200 | 199.0 | 17.0 | 217 | 22 | 133 | 9,07E+07 | 3 | 404 | 0 | 99 | 0 | 1 |
| ZA17 | 29 July 2014 | 7 | 31 | 9.7 | 12600 | 6.8 | 10.1 | 142 | 8 | 502 | 6,20E+07 | 144 | 62 | 0 | 26 | 397 | 419 |

n.d. - no data
<sup>1</sup>POC = TOC - DOC; ISS = TSS - POC
<sup>2</sup>CyP - cyanobacterial plankton
<sup>3</sup>EuAPP - eukaryotic autotrophic picoplankton (= pico-sized eukaryotic algae)

**Supplementary Table S2.**
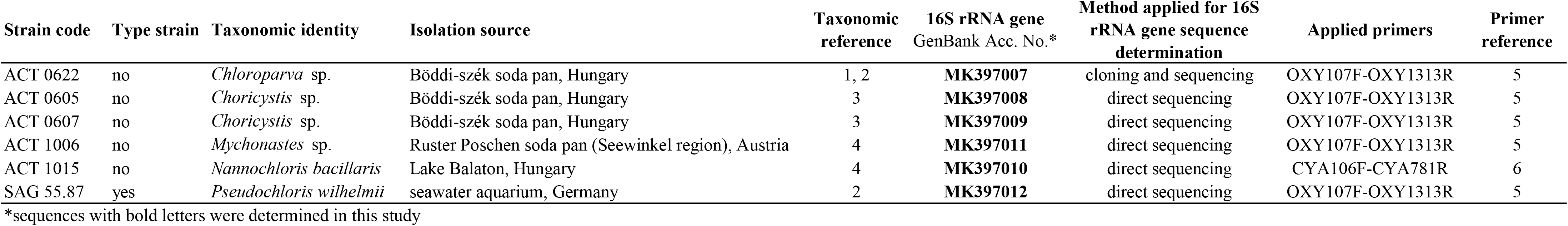
Reference chloroplast 16S rRNA gene sequences generated in this study

**Supplementary Table S3.** Eukaryotic nanoplanktonic algae detected in the pans and their estimated biomass

**turbid pan (Zab-szék)**
| Sampling date | 17 April 2013 | 30 May 2013 | 18 June 2013 | 24 July 2013 | 16 August 2013 | 24 September 2013 | 17 October 2013 | 19 November 2013 |
| --- | --- | --- | --- | --- | --- | --- | --- | --- |
| *Cryptomonas* sp. (µg l<sup>-1</sup>) | 0 | 0 | 0 | 0 | 0 | 0 | 0 | 0 |
| *Chlamydomonas* sp. (µg l<sup>-1</sup>) | 0 | 0 | 0 | 0 | 27 | 0 | 0 | 0 |
| *Nitzschia palea* (µg l<sup>-1</sup>) | 0 | 0 | 0 | 0 | 12 | 0 | 0 | 0 |
| Sum of the biomass of eukaryotic nanoplanktonic algae (µg l<sup>-1</sup>) | 0 | 0 | 0 | 0 | 39 | 0 | 0 | 0 |

**colored pan (Sós-ér)**
| Sampling date | 17 April 2013 | 30 May 2013 | 18 June 2013 | 24 July 2013 | 16 August 2013 | 24 September 2013 | 17 October 2013 | 19 November 2013 |
| --- | --- | --- | --- | --- | --- | --- | --- | --- |
| *Euglena* sp. (µg l<sup>-1</sup>) | 0 | 0 | 0 | 0 | 4000 | n.a. | n.a. | 520 |
| *Phacus* sp. (µg l<sup>-1</sup>) | 0 | 0 | 0 | 0 | 0 | n.a. | n.a. | 0 |
| *Cryptomonas* sp. (µg l<sup>-1</sup>) | 6 | 0 | 0 | 0 | 0 | n.a. | n.a. | 0 |
| *Rhodomonas* sp. (µg l<sup>-1</sup>) | 3 | 0 | 4 | 0 | 0 | n.a. | n.a. | 0 |
| *Tribonema* sp. (µg l<sup>-1</sup>) | 0 | 150 | 0 | 0 | 0 | n.a. | n.a. | 0 |
| *Carteria* sp. (µg l<sup>-1</sup>) | 0 | 0 | 0 | 0 | 0 | n.a. | n.a. | 2040 |
| *Chlamydomonas* sp. (µg l<sup>-1</sup>) | 5 | 0 | 0 | 0 | 28150 | n.a. | n.a. | 3400 |
| *Nitzschia* sp. (µg l<sup>-1</sup>) | 0 | 2 | 0 | 10 | 0 | n.a. | n.a. | 0 |
| *Navicula* sp. (µg l<sup>-1</sup>) | 0 | 0 | 0 | 0 | 0 | n.a. | n.a. | 0 |
| *Cocconeis* sp. (µg l<sup>-1</sup>) | 0 | 0 | 0 | 12 | 0 | n.a. | n.a. | 0 |
| *Cymbella* sp. (µg l<sup>-1</sup>) | 0 | 0 | 0 | 13 | 0 | n.a. | n.a. | 0 |
| Sum of the biomass of eukaryotic nanoplanktonic algae (µg l<sup>-1</sup>) | 14 | 152 | 4 | 35 | 32150 | n.a. | n.a. | 5960 |

| 09 December 2013 | 10 January 2014 | 26 February 2014 | 26 March 2014 | 23 April 2014 | 22 May 2014 | 18 June 2014 | 29 July 2014 |
| --- | --- | --- | --- | --- | --- | --- | --- |
| 0 | 0 | 0 | 3450 | 0 | 0 | 0 | 0 |
| 0 | 0 | 0 | 0 | 0 | 0 | 0 | 0 |
| 40 | 0 | 0 | 0 | 0 | 0 | 0 | 0 |
| 40 | 0 | 0 | 3450 | 0 | 0 | 0 | 0 |
| 2340 | 8250 | 3960 | 9600 | 0 | 0 | 1200 | n.a. |
| 0 | 0 | 0 | 2240 | 0 | 0 | 500 | n.a. |
| 0 | 0 | 0 | 0 | 0 | 0 | 0 | n.a. |
| 0 | 0 | 0 | 0 | 0 | 0 | 0 | n.a. |
| 0 | 0 | 0 | 0 | 0 | 0 | 0 | n.a. |
| 600 | 0 | 0 | 0 | 0 | 0 | 0 | n.a. |
| 40 | 3420 | 4200 | 0 | 75 | 0 | 30 | n.a. |
| 8 | 0 | 0 | 0 | 0 | 0 | 0 | n.a. |
| 0 | 0 | 0 | 0 | 0 | 0 | 980 | n.a. |
| 0 | 0 | 0 | 0 | 0 | 0 | 0 | n.a. |
| 0 | 0 | 0 | 0 | 0 | 0 | 0 | n.a. |
| 2988 | 11670 | 8160 | 11840 | 75 | 0 | 2710 | n.a. |

